# PARP14 inhibits the inflammatory response of macrophages through the NF-κB pathway

**DOI:** 10.1101/2024.03.12.584632

**Authors:** Xuefei Guo, Yang Zhao

## Abstract

The poly-ADP-ribose polymerase (PARP) superfamily consists of 17 members, which regulate many biological processes in physiological or pathological conditions, such as DNA damage repair, anti-viral responses, and development of adaptive immune cells. Among them, PARP14 is the biggest member, containing two RNA recognition motifs at the N-terminal, three macro-domains, one WWE domain, and one PARP domain at the C-terminal, which was reported to regulate IL4/STAT6 signaling in adaptive immune cells. However, whether PARP14 participates in regulating host inflammatory response remains unclear. In a previous study, we observed that virus infection and LPS treatment induced the transcription of *Parp14*. By comparing the primary macrophages derived from *Parp14* KO and WT mice, we found that some inflammatory cytokines were significantly induced in KO macrophages. Still, the expression of *Ifnb1* had no significant difference compared to the WT macrophages. RNA-seq analysis showed that the KO group had a more robust inflammatory response but a weaker innate immune response upon stimulation. We validated these results by performing a knockdown of *Parp14* in RAW 264.7 cells. Moreover, the survival time of the KO mice was much shorter than that of the WT group upon LPS injection. Transcription factor enrichment analysis indicated that nuclear factor-kappaB1 (NF-κB1) may be the main reason for increasing the production of these inflammatory cytokines. As expected, the up-regulation was deleted upon the treatment of the inhibitor of NF-κB, JSH23. These data imply that PARP14 regulates inflammatory responses through the NF-κB pathway.

## Introduction

Inflammation is a fundamental and complex biological response triggered by various harmful stimuli, including infections, injuries, and toxins, which functions as a defense mechanism at an early stage^1,2^. While acute inflammation serves to protect the body, chronic inflammation, often driven by environmental factors like smoking, obesity, and stress, can contribute to kinds of diseases containing atherosclerosis, rheumatoid arthritis, and cancer^3^. The immune system may remain activated over extended periods, leading to tissue damage and dysfunction in the chronic inflammatory conditions^4^. Understanding the intricate mechanisms underlying inflammation is critical for developing targeted therapies for inflammatory diseases. Previous studies showed that NF-κB is central in orchestrating the inflammatory response^5^. Its intricate signaling pathways are essential for the host’s defense against pathogens and injury, but when dysregulated, they can contribute to chronic inflammatory conditions^6^. Although increasing studies continue to uncover the regulation of inflammation, the precise regulatory mechanisms of NF-κB remain to be uncovered.

The poly-ADP-ribose polymerase (PARP) family is a group of enzymes with diverse functions, which have garnered increasing attention due to their critical roles in various cellular processes in recent years, including responding to cellular stress, DNA damage and repair, regulating gene expression, and cell death^7–9^. PARP enzymes could catalyze the transfer process of ADP-ribose units from nicotinamide adenine dinucleotide (NAD^+^) to target proteins, the process also known as ADP-ribosylation^10^. This modification could change the structure and function of target proteins, impacting their involvement in numerous cellular pathways. The PARP superfamily contains 17 members in humans, each with distinct functions and specificities^7^. One of the well-known members of this superfamily is PARP1, primarily involved in DNA repair and promoting the stability of the genome^11^. PARP2, a close relative of PARP1, shares many functions, although their specific roles can vary in cellular contexts^12^. Furthermore, PARP3 and PARP16 also have roles in DNA repair and other cellular processes, but their functions could be more well characterized^13,14^. Among them, PARP14 was the biggest protein, containing 2 RNA-binding motifs, 3 macrodomains, 1 WWE domain, and 1 PARP domain^15–17^. The reported functions of PARP14 include participating in the DNA damage and repair, binding the promoter of STAT6 upon the IL4 treatments, then inducing the production of downstream cytokines, and promoting the survival of multiple myeloma cell lines by the JNK signaling^18–25^. Our previous study found that the transcription of *Parp14* could be induced significantly by viral infection or LPS treatment. However, the roles of PARP14 in inflammatory response are still unclear.

In this study, we separated the primary cells from *Parp14* knockout (KO) and wild-type (WT) mice, such as bone marrow-derived macrophages (BMDMs), peritoneal macrophages (PMs), and spleen-derived lymphocytes. After treatment with VSV, HSV, MHV, SeV, or LPS, we found that the production of type I interferon IFNB1 by murine-derived primary cells in the KO group was weaker than in the WT group. However, the induction of inflammatory cytokines was much stronger than the WT group. To further clarify the biological processes and signaling pathways involved in PARP14, we used PMs and BMDMs to conduct the RNA-seq. Subsequently, we used shRNA to knock down *Parp14* in RAW 264.7 cells to validate these results. Then, we constructed a sepsis model using LPS to demonstrate the anti-inflammatory effect of PARP14 at the animal level. By performing the transcription factor enrichment analysis, NF-κB1 and IRF8 were identified as the potential factors leading to the difference. Finally, we found that PARP14 inhibits the expression of some inflammatory cytokines in macrophages through the NF-κB pathway.

## Materials and methods

### Cell culture and viral infection

GM-CSF cytokines induced the bone marrow-derived cells isolated from male C57BL/6J mice for seven days. We keep the cells with DMEM containing 10% FBS during the induction. On day 7, we harvested the Bone Marrow-Derived Macrophages (BMDMs) to a 12-well plate and infected them with SeV, MHV, HSV, VSV, or other stimuli for 6 hours, respectively. The RAW 264.7 cells were also cultured with DMEM as BMDMs at the 37°C CO2 (5%) incubator. Incubate the cells until they reach the desired confluence or are ready for experimental procedures. Check cells daily for confluence and media conditions.

### Quantitative RT-PCR

The TRIZol reagent (TIANGEN) was used to extract the total RNA from cells. The purified RNA was reversely transcribed using HiScript II RT SuperMix (Vazyme). Then, the expression level of target genes was counted by SYBR Green qMix (Vazyme). Data of RT-qPCT shown in this study were the relative expression level of targeted mRNA standardized to the *Actb*. All the primers used in the present study were listed in the supplementary table: Table S3.

### CRISPR/Cas9 to knock out *Parp14*

The Clustered Regularly Interspaced Short Palindromic Repeats (CRISPR) technology is a powerful method for genome editing, allowing precise modification of DNA sequences in cells or organisms^36^. The *Parp14* KO mice were bought from the Cyagen company (https://www.cyagen.com/cn/zh-cn/). The primer sequences of CRISPR-Cas9 to generate *Parp14* KO mice and the coordinated validated primer sequences are shown in the supplementary table: Table S1.

### shRNA to knock down *Parp14*

The Short Hairpin RNA (shRNA) technology is widely used for silencing specific target genes in cells. The present study used the pLKO.1 plasmid as a vector containing EcoR1 and AgeI restriction enzyme cutting sites. The shRNA sequences to knock down *Parp14* were designed with this online database (https://www.sigmaaldrich.cn/CN/zh/product/sigma/shrna), as shown in the supplementary table: Table S2. We designed two pairs of shRNA for *Parp14* first. After validation, we chose the second pair for further experiments because of a much higher knockdown efficiency for *Parp14*.

### RNA-seq sample preparations

Upon HSV and VSV infection or LPS treatment for 6 hours, we obtained the total RNA from BMDMs or PMs using the TRIZol reagent (TIANGEN). The purified were used to construct the RNA libraries, then sequenced by the GENEWIZ company (https://www.genewiz.com/). After the sequencing is completed, the company will send the raw data in fasta file format.

### RNA-seq data analysis

The FastQC and Trim-Galore software were used to quality control raw data to generate clean data. Then, the clean data was aligned to their reference genome, such as hg19, rheMac10, and mm10, with Subread software^37^, respectively. The software feature Counts from Subread was used to obtain the gene expression matrix^38^. Then, we used the FPKM formula to normalize the coding gene expression matrix. The R package DESeq2 was used for the differentially expressed analysis to identify the differentially expressed genes (DEGs)^39^.

### GO annotation and KEGG pathway enrichment analysis

In this study, these identified DEGs were enriched by the GO annotation and KEGG pathway enrichment analysis with the clusterProfiler R package (the enrichGO and enrichKEGG functions, respectively)^40^ or the online database DAVID^41^. All the GO and KEGG terms with *FDR* less than 0.01 were considered significant enrichments.

### Gene set enrichment analysis (GSEA)

The GSEA is an important computational way to determine if predefined sets of genes exhibit statistically significant and concordant differences between two biological states, whose input is all the genes but not only those DEGs meeting the significance level. The clusterProfiler package was also utilized to conduct the GSEA in this study. Then, the function gseaplot2 from clusterProfiler was used to visualize the enrichment analysis results.

### Protein-protein interaction (PPI) network analysis

PPI network analysis is a computational and bioinformatics approach used to explore and understand the interactions among proteins in a biological system^42^. The online database (https://string-db.org/) was used to perform this analysis with default settings, whose input is the DEGs and output is the network between the proteins of DEGs. Then, the software Cytoscape was utilized for visualization^43^.

### Transcription factor enrichment analysis (TFEA)

TFEA is a bioinformatics approach to identify transcription factors (TFs) likely to regulate differentially expressed genes. iRegulon (http://iregulon.aertslab.org/tutorial.html) is a plug-in of Cytoscape which was designed for conducting TFEA, whose input was a list of DEGs, whose output was a list of predicted TFs with a rank of enrichment score^44^. The results of TFEA could be visualized by the Cytoscape software directly^45^.

### LPS and VSV injection

LPS-induced sepsis models are commonly used to study the systemic inflammatory response seen in sepsis. In the present study, we dissolved the LPS in sterile saline to the desired concentration (20 mg/kg for mice). Administer the LPS via the intraperitoneal (IP) route using a sterile insulin syringe. Record the LPS concentration, the injection volume, and administration time. Record vital signs as time passes, including body temperature and behavioral changes. Unlike the injection of LPS, we infected the mice with VSV using the way of the tail vein (1×10^8 plaque-forming units per milliliter in this study). After injection, record the survival time of mice. We collected blood, tissues, and other organs for further analysis.

### Statistical analysis

In the present study, the Python package SciPy (https://pypi.org/project/scipy/) was used to conduct all the statistical analyses as indicated. Results are presented as the mean ± SEM. The *p* values < 0.05 were considered statistically significant (*), *p* values < 0.01, and *p* values < 0.001 were regarded as highly statistically significant (** and ***).

## Results

### Viral infection up-regulated the expression of PARP14 in macrophages

In the previous study, we observed that most of the PARP members could be up-regulated upon both VSV and HSV infection, such as *Parp3*, *Parp8*, *Parp9*, *Parp10*, *Parp11*, *Parp12*, *Parp13*, *Parp14,* and so on, which correlated with the expression of *Ifnb1*, *Isg15*, and *Cxcl10* (Fig 1A). As shown in Fig 1B, we found that *Parp14* was also up-regulated in the lung tissue of *Rhesus macaques* infected by SARS-CoV-2 on day 3 and day 7 through re-analyzing the publicly published data (GSE158297). These data indicated that *Parp14* may play a role in the anti-viral process of the host. Then, we search for the expression pattern of *PARP14* in the Human Protein Atlas database (https://www.proteinatlas.org/)^26^. Data showed that *Parp14* was widely expressed in different human tissues; the spleen, tonsil, liver, liver, lung, and so on were the top 10 tissues (Fig 1C). It was also worth noting that the *Parp14* gene was highly expressed in the human immune cells, such as B-cells, Macrophages, Monocytes, NK cells, and T cells (Fig 1D), suggesting its function may relate to the host’s immune system. To validate these predictions, we examined the expression of Parp14 at different immune cells with various viruses infection (Fig 1E-G). The results shown in Fig 1E show that MHV, HSV, and VSV infection could significantly induce the expression of Parp14 compared to the mock group in BMDMs, except for the SeV infection. In the PMs and RAW 264.7 cells, all of these viruses up-regulated the expression level of *Parp14* significantly (Fig 1F-G). Together, these results demonstrated that virus infection could up-regulate the expression of *Parp14* in macrophages.

**Fig. 1.**
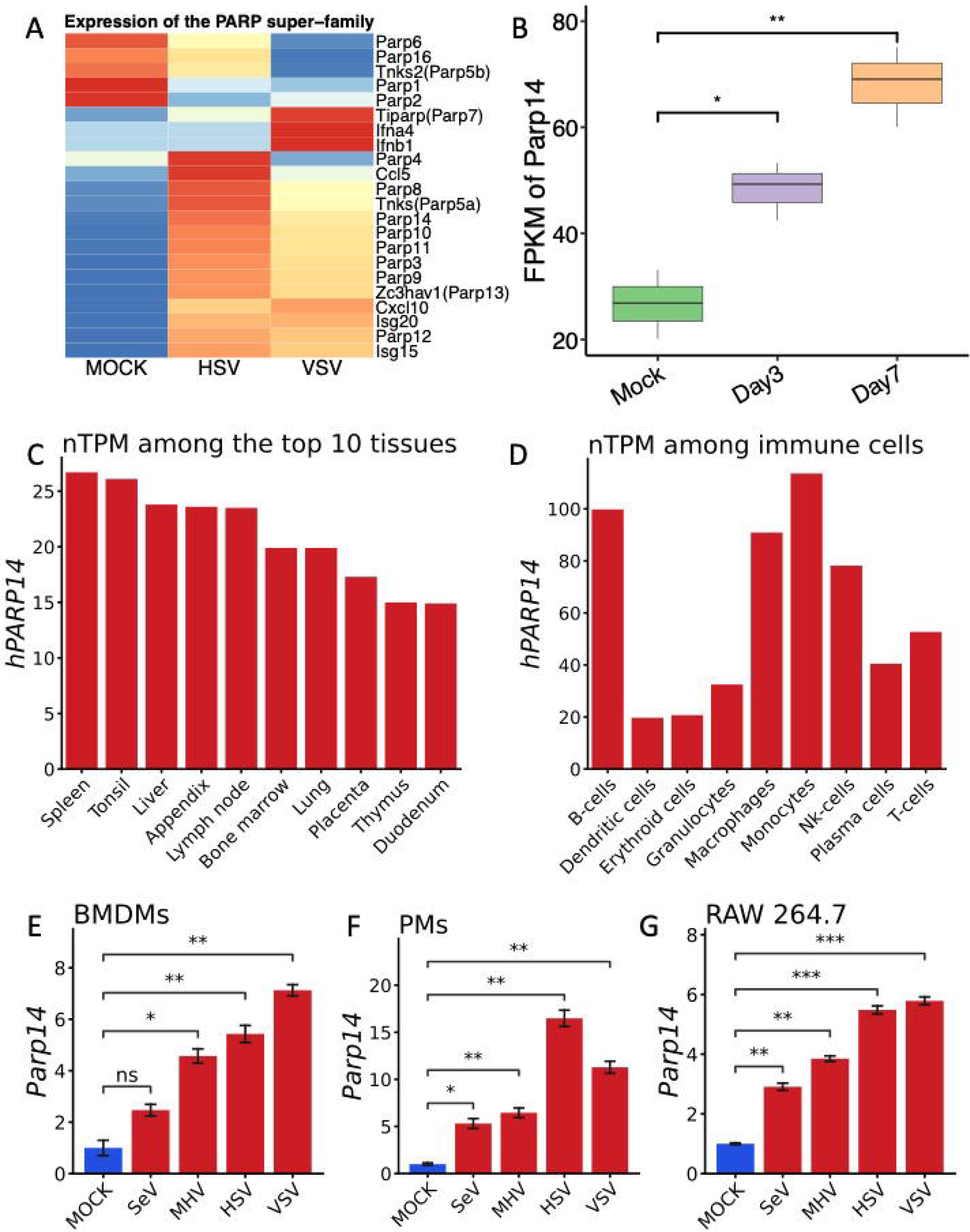
Virus infection up-regulated the expression of *Parp14* in macrophages. A) HSV and VSV infection up-regulated most members of the PARP superfamily. B) SARS-CoV-2 infection up-regulated the transcription of *Parp14* in the lung tissue of *Rhesus macaques*. C) The top 10 human tissues highly expressed the *PARP14*. D) The top 9 human immune cells highly expressed the *PARP14*. E-G) Various virus infections up-regulated the expression of *Parp14* at BMDMs, PMs, and RAW 264.7 cells, respectively.

### Depletion of PARP14 enhances the expression of *Il1b* and *Il6* in macrophages

Then, we generated the *Parp14* knock-out (KO) mice and macrophages to explore the potential roles of *Parp14* in the host immune system. Unexpectedly, the transcription of *Ifnb1* from the WT and KO groups had no significant difference with the infection of HSV or VSV for 6 hours (Fig 2A-B). However, the expression of *Il1b* and *Il6* were significantly up-regulated in *Parp14* KO PMs upon the infection of HSV and VSV, respectively (Fig 2A-B). Consistent with the results of PMs, the expression of *Ifnb1* in the *Parp14* KO was weaker than WT BMDMs but did not meet the significant difference upon the infection of HSV, SeV, and VSV (Fig 2C). In contrast, the *Il6* had a more robust expression in *Parp14* KO BMDMs upon viral infection.

**Fig. 2.**
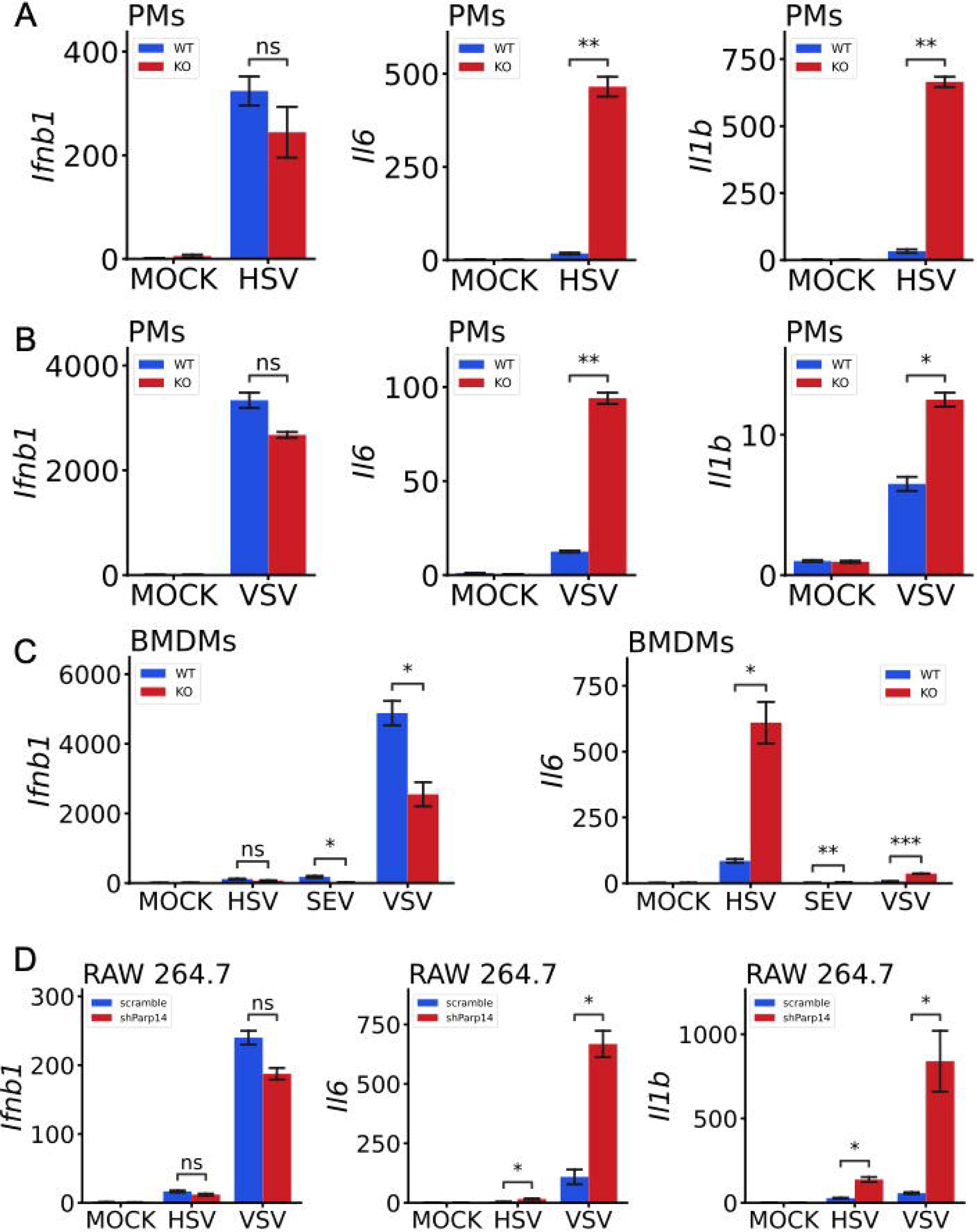
Depletion of *Parp14* enhanced the transcription of *Il-1b* and *Il-6* in macrophages. A) The expression of *Ifnb1*, *Il6*, and *Il1b* at *Parp14* KO and WT PMs upon HSV infection. B) The expression of *Ifnb1*, *Il6*, and *Il1b* at *Parp14* KO and WT PMs upon VSV infection. C) The expression of *Ifnb1* and *Il6* at *Parp14* KO and WT BMDMs upon HSV, SEV, and VSV infection. D) The expression of *Ifnb1*, *Il6*, and *Il1b* at *Parp14* KD and WT RAW 264.7 cells upon HSV and VSV infection.

To validate these results, we used the shRNA of *Parp14* to generate the *Parp14* knock-down (KD) RAW 264.7 cells. The primer sequences of shRNA for *Parp14* are shown in Table S2. The effect of shRNA to inhibit the expression of *Parp14* was shown in Fig S1A, which could significantly down-regulate the expression of *Parp14* at RAW 264.7 cells with or without virus treatment. Consistent with these results from BMDMs and PMs, the expression of *Il1b* and *Il6* rather than *Ifnb1* were significantly up-regulated in KD RAW 264.7 cells upon HSV and VSV treatment compared to the WT RAW 264.7 cells. We also found that the expression of *Cxcl2* was up-regulated considerably upon HSV and VSV treatment (Fig S1B). Overall, these results demonstrated that KO or KD *Parp14* in the macrophages almost had no significant influence on *Ifnb1* but significantly elicited the expression of *Il1b* and *Il6*, which indicated that *Parp14* may play roles in the host’s inflammatory response upon stimulations.

### KO of PARP14 exhibits a more robust inflammatory response upon virus infection in macrophages

To depict the influence of *Parp14* KO on host cells at the whole transcriptome level, we conducted the bulk RNA-seq on *Parp14* KO and WT BMDMs upon HSV and VSV infection. As shown in Fig 3A, some genes were up-regulated or down-regulated regardless of HSV or VSV infection. For example, *Arg1*, *Csta2*, *Csf3*, *Il1b*, *Pf4*, and *S100a8* genes were induced more highly upon both HSV and VSV infection in the KO group compared to the WT group, while *Tlr9*, *Wdfy1,* and *Itpr1* genes in KO group were lower than WT BMDMs. Fig 3B shows that *Parp14* and some interferon genes were significantly induced in WT BMDMs upon HSV and VSV treatment. At the same time, some inflammatory cytokines, such as *Il1a*, *Il1b*, *Pf4*, *Cxcl2*, and *Tnfsf9,* had a higher expression level in the KO BMDMs group. Then, we performed the KEGG pathway enrichment and GO annotation analysis for the DEGs between the KO and WT BMDMs upon HSV and VSV treatment, respectively (Fig 3C-F). As shown in Fig 3C and Fig 3E, the up-regulated genes were enriched into these pathways, which included Hematopoietic cell lineage, Rheumatoid arthritis, Cytokine-cytokine receptor interaction, Th17 cell differentiation, Th1 and Th2 cell differentiation, T cell receptor signaling pathway and inflammatory bowel disease and so on regardless of HSV or VSV infection. The GO annotation results of up-regulated DEGs upon HSV and VSV treatment were shown in Fig 3D and Fig 3F: immune system process, adaptive immune response, MHC class II protein complex, positive regulation of T cell activation, cellular response to interferon-pathway and neutrophil chemotaxis.

**Fig. 3.**
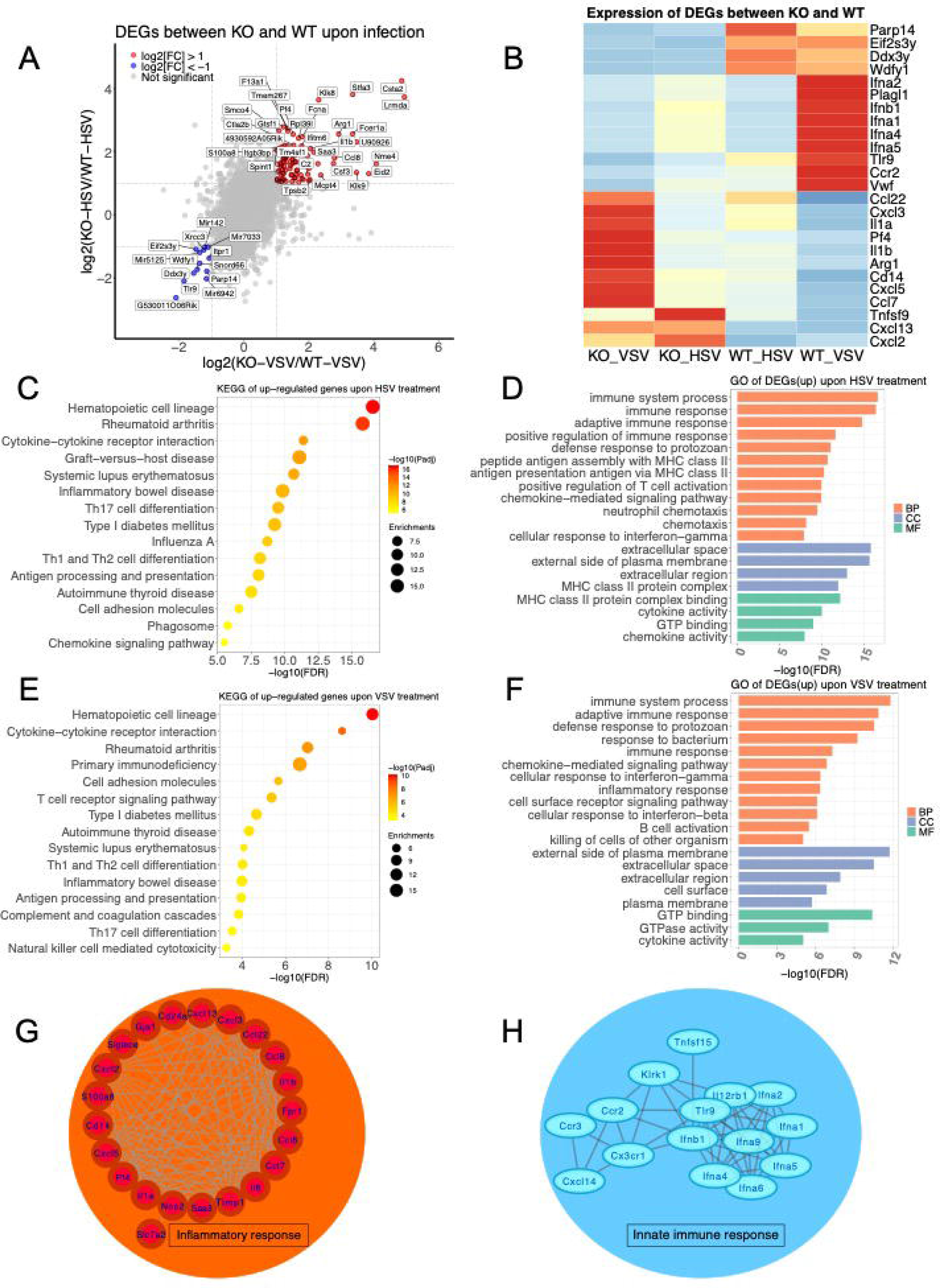
*Parp14* KO macrophages exhibited a more robust inflammatory response upon virus infection. A) The dot plot representing DEGs upon HSV and VSV infection between *Parp14* KO and WT BMDMs. B) The heatmap of DEGs between the *Parp14* KO and WT BMDMs upon HSV and VSV infection. C-D) The KEGG and GO enrichment analysis of up-regulated DEGs in the *Parp14* KO group compared to the WT BMDMs upon HSV treatment. E-F) The KEGG and GO enrichment analysis of up-regulated DEGs in the *Parp14* KO group compared to the WT BMDMs upon VSV treatment. G) The PPI network analysis of up-regulated DEGs in the *Parp14* KO group compared to the WT BMDMs upon VSV treatment. H) The PPI network analysis of down-regulated DEGs in the *Parp14* KO group compared to the WT BMDMs upon VSV treatment.

The GO enrichment analysis results of down-regulated DEGs in the KO BMDMs were shown in Fig S2A-B, which contained the protein monoubiquitination, gene silencing by RNA, regulation of phosphorylation of STAT proteins, and immune response. The KEGG enrichment analysis of significantly down-regulated genes in the KO group upon VSV treatment included some innate immune-related responses, such as NOD-like receptor signaling pathway, Toll-like receptor signaling pathway, RIG-I-like receptor signaling pathway, IL-17 signaling pathway (Fig S3C). Then, we focused on the DEGs between *Parp14* KO and WT upon VSV infection by conducting a PPI network analysis. As shown in Fig 3G-H, some up-regulated DEGs were enriched in the inflammatory response, while the down-regulated DEGs were enriched in the innate immune response. Together, these results suggested that KO of *Parp14* impaired the innate immune response but enhanced the inflammatory response of the host.

### Knockout of Parp14 increases the susceptibility to LPS challenge

Because the *Parp14* KO macrophages exhibited a more robust inflammatory response, we examined whether there was a distinguished response to the LPS challenge between the KO and WT groups. As shown in Fig 4A, LPS treatment could not generate a significant difference in the expression of *Ifnb1* but significantly induced the transcription of *Il6* and *Il1b*, consistent with the treatment of HSV and VSV. Moreover, LPS treatment also induced a higher expression level of inflammatory cytokines rather than the expression of *Ifnb1* (Fig 4B). To further validate the anti-inflammatory effect of *Parp14*, we injected the *Parp14* KO and WT mice with VSV and LPS, respectively. As expected, the survival time of KO and WT mice had no significant difference upon VSV infection, consistent with these results at the cell level. However, the survival time of WT mice was significantly longer than the *Parp14* KO mice. Then, we examined the expression of some inflammatory cytokines between the KO and WT mice. As shown in Fig 4E, upon LPS injection 10 hours, the *Tnf*, *Il6,* and *Il1b* genes were significantly higher than the WT mice, which may be the main reason for the death of *Parp14* KO mice. Together, these data demonstrated that *Parp14* played a stronger anti-inflammatory function than its anti-viral effect.

**Fig. 4.**
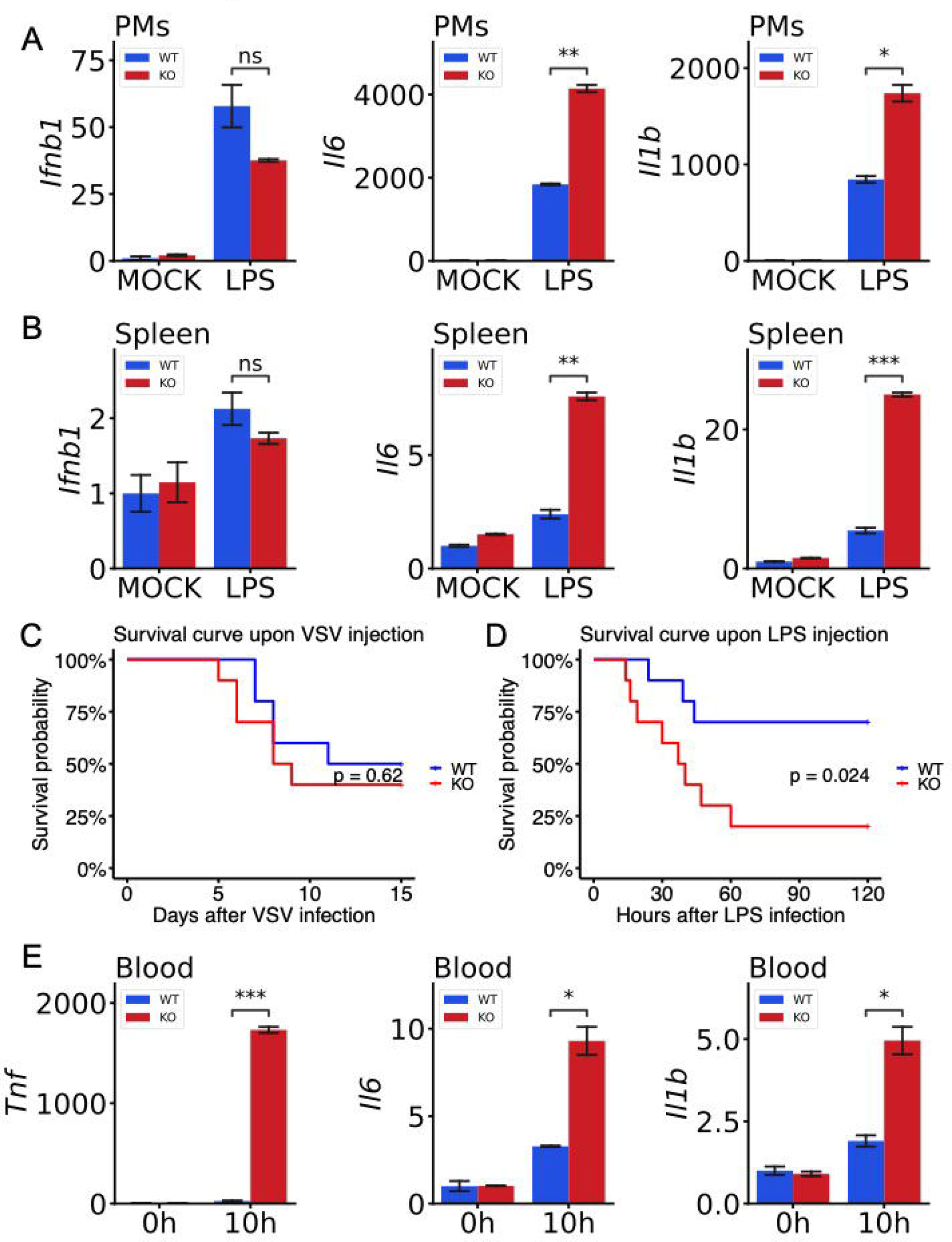
Knock-out of *Parp14* increases the susceptibility to LPS challenge. A) The expression of *Ifnb1*, *Il6*, and *Il1b* in the PMs primary cell upon LPS treatment. B The expression of *Ifnb1*, *Il6*, and *Il1b* in the spleen primary cell upon LPS treatment. C-D) The survival curve of *Parp14* KO and WT mice upon VSV and LPS injection, respectively. E) The expression of *Tnf*, *Il6,* and *Il1b* of the peripheral blood from KO and WT mice upon LPS injection for 10 hours.

### PARP14 inhibits the inflammatory response through the NKFB pathway

To further explore the molecular mechanism by which *Parp14* plays the anti-inflammatory role, we conducted the RNA-seq on the PMs derived from the KO and WT mice and then treated them with LPS for 6 hours. The DEGs shown in Fig 5A, *Arg1*, *Cxcl13*, *Gzma*, *Gzmc*, *Il1b,* and *Cxcr3* were significantly up-regulated, while *Fgg*, *Fga*, *Fgb*, *Apoa1*, *Apoh,* and *Alb* were down-regulated considerably in KO compared to WT group. KEGG pathway enrichment analysis was conducted on the up-regulated DEGs in the KO group, and the results were shown in Fig 5B, which contained adaptive immune response, leukocyte cell-cell adhesion, regulation of T cell activation, leukocyte migration, lymphocyte differentiation and lymphocyte-mediated immunity and so on. Then, we conducted the GSEA between KO and WT PMs (Fig 5C-D). The top enriched terms included regulation of innate immune response, positive regulation of inflammatory response, neutrophil chemotaxis, neutrophil migration, leukocyte chemotaxis, adaptive immune response, positive regulation of lymphocyte activation, and lymphocyte-mediated immunity (Fig S3A-F).

**Fig. 5.**
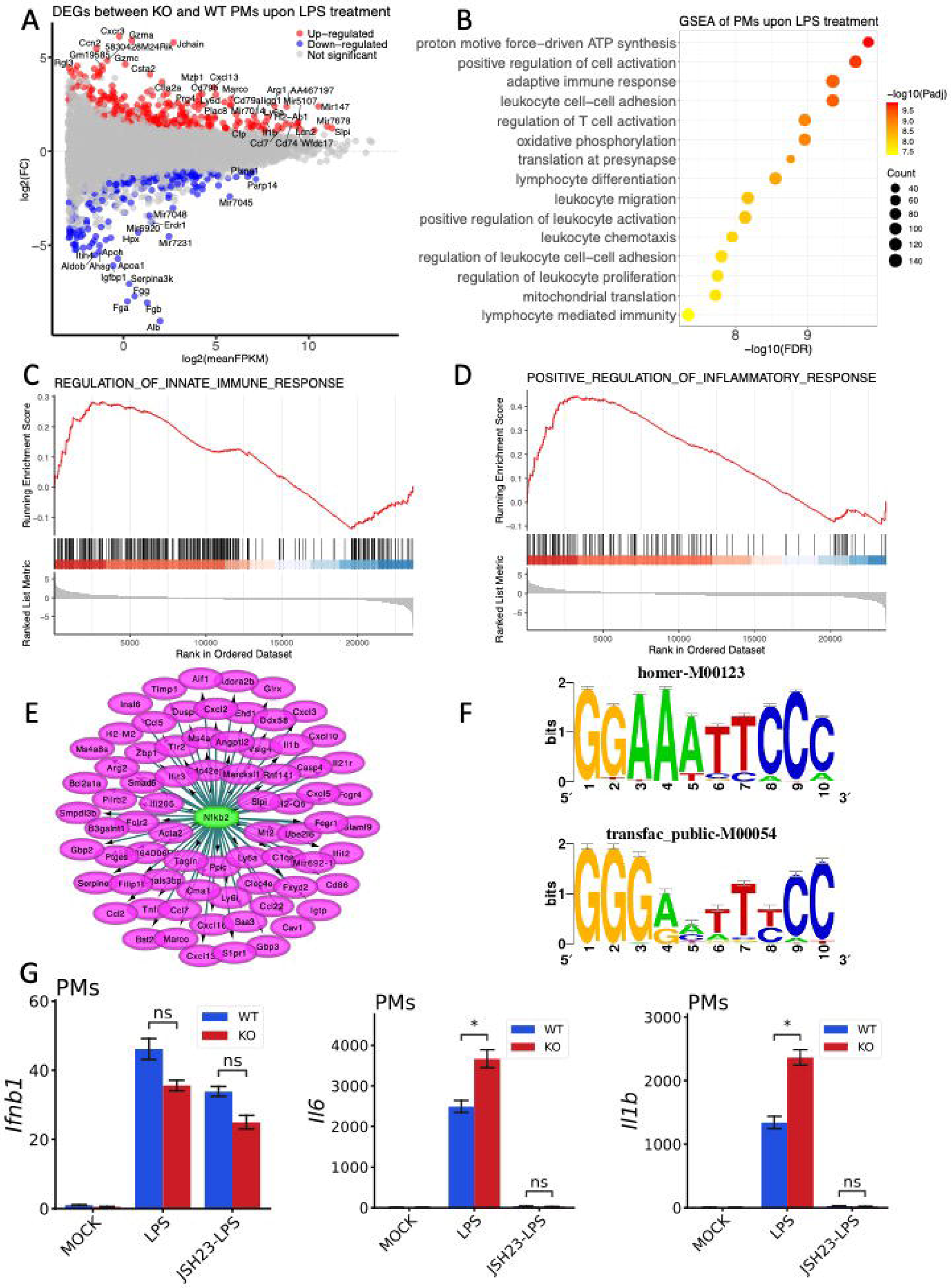
*Parp14* inhibits the inflammatory response through the NF-κB signaling pathway. A) The MA plot of DEGs between KO and WT PMs upon LPS treatment. B) The KEGG enrichment analysis of up-regulated DEGs in the KO group compared to the WT PMs upon LPS treatment. C-D) The GSEA enrichment analysis of DEGs between KO and WT PMs upon LPS treatment. E) The transcription factor enrichment analysis (TFEA) of up-regulated genes in the KO group compared to the WT PMs upon LPS treatment. F) The predicted DNA motif sequences of NF-κB1 to bind. G) The NF-κB inhibitor, JSH23, eliminated the difference of *Il6* and *Il1b* between WT and KO PMs.

Then, we used the up-regulated DEGs in KO PMs to perform the TFEA with Cytoscape software and iRegulon plug-in. As shown in Fig 5E, NF-κB1 was the top enriched transcription factor (TF) that regulated most of these up-regulated genes. The predicted DNA motif sequences bound by the NF-κB1 are shown in Fig 5F. The second enriched TF was IRF8; the predicted binding motifs are shown in Fig S4A-B. These results indicated that *Parp14* may regulate the expression of inflammatory cytokines through the NF-κB or IRF8. Thus, we used the inhibitor of NF-ΚB, JSH23, to pre-treat KO and WT PMs before LPS treatment. As shown in Fig 5G, JSH23 almost eliminated the difference of *Il6* and *Il1b* between WT and KO groups, which demonstrated that *Parp14* inhibited the expression of some inflammatory cytokines through the NF-κB pathway.

## Discussion

There are 17 members in the PARP superfamily being found. Previous studies reported that some members of the PARP family play crucial roles under various physiological or pathological conditions. Among them, PARP1 and PARP2 were studied well, participating in the DNA damage and repair process^11–13^. PARP13 was also a well-established ISG that could inhibit the replication of PARP13 directly to defend against the infection of viruses^27^. The other members of the PARP superfamily are less well-known. PARP14 is the biggest member of the PARP superfamily, with more than 1800 amino acids, containing 2 RNA-binding motifs, 3 macro-domains, 1 WWE domain, and 1 PARP domain^28^. The reported functions of PARP14 contain participating in the DNA damage and repair, promoting the stability of the genome, binding the promoter of STAT6 upon the IL4 treatments, then inducing the production of downstream cytokines, promoting the survival of multiple myeloma cell lines by the JNK signaling^15,19,20,29,30^. However, the detailed roles of PARP14 in inflammation are still unclear.

This study compared the primary cells from *Parp14* KO and WT mice, such as PMs, BMDMs, and spleen-derived lymphocytes. Furthermore, it found that after treatment by VSV, HSV, MHV, SeV, or LPS, some inflammatory cytokines were more potent than those in the WT group, but the production of *Ifnb1* had no significant difference. To further clarify the biological processes and signaling pathways involved in PARP14, we used PMs and BMDMs to perform the RNA-seq. The analysis results confirmed that the KO group had a more robust inflammatory response but a weaker innate immune response upon stimulation. Subsequently, we used shRNA to knock down *Parp14*, whose results were consistent with the primary macrophages derived from *Parp14* KO and WT mice. Moreover, we constructed a sepsis model using LPS to verify the anti-inflammatory effect of *Parp14* at the animal level. The survival time of mice in the KO group was much lower than that in the WT group due to the high expression of various inflammatory cytokines, resulting in a stronger cytokine storm induced by LPS. TFEA results indicated that NF-κB1 and IRF8 may lead to the up-regulation of these inflammatory cytokines. As expected, the up-regulation was deleted upon treating the inhibitor of NF-ΚB, JSH23. However, how PARP14 interacts with NFKB and the roles of IRF8 in the inflammatory response need much work to clarify^31^.

Although the RNA-seq results showed that the *Parp14* KO group exhibited a weaker innate immune response than the WT group (Fig 3H), there was no significant difference in survival time between the WT and KO groups after the VSV infection. Moreover, the expression level of some ISGs in the KO and WT groups was almost comparable upon VSV injection, such as *Isg15* and *Cxcl10*. One potential reason for this phenomenon was that the baseline of these ISGs in the KO was much higher than in the WT group without any treatment. This was because PARP14 could inhibit the phosphorylation of STAT1^30^, which positively regulates the expression of some ISGs. After VSV or HSV treatment, the expression of *Ifnb1* was much higher in the WT group, which could induce a higher level of downstream ISGs than in the KO group. Consequently, there was almost no difference in the expression of ISG levels between the KO group and the WT group upon VSV infection. However, whether PARP14 KO influences the function of pattern recognition receptors, such as MDA5, RIG-I, and cGAS, needs more experiments to demonstrate in future studies^32–35^. Overall, these data indicate the role of PARP14 in controlling inflammatory responses through the NF-κB pathway upon viral infection or LPS treatment.

## Acknowledgments

We thank the support from the High-performance Computing Platform of Peking University for providing the computing clusters to facilitate the data analysis procedure in this research.

## Supplementary Materials

**Fig. S1.**
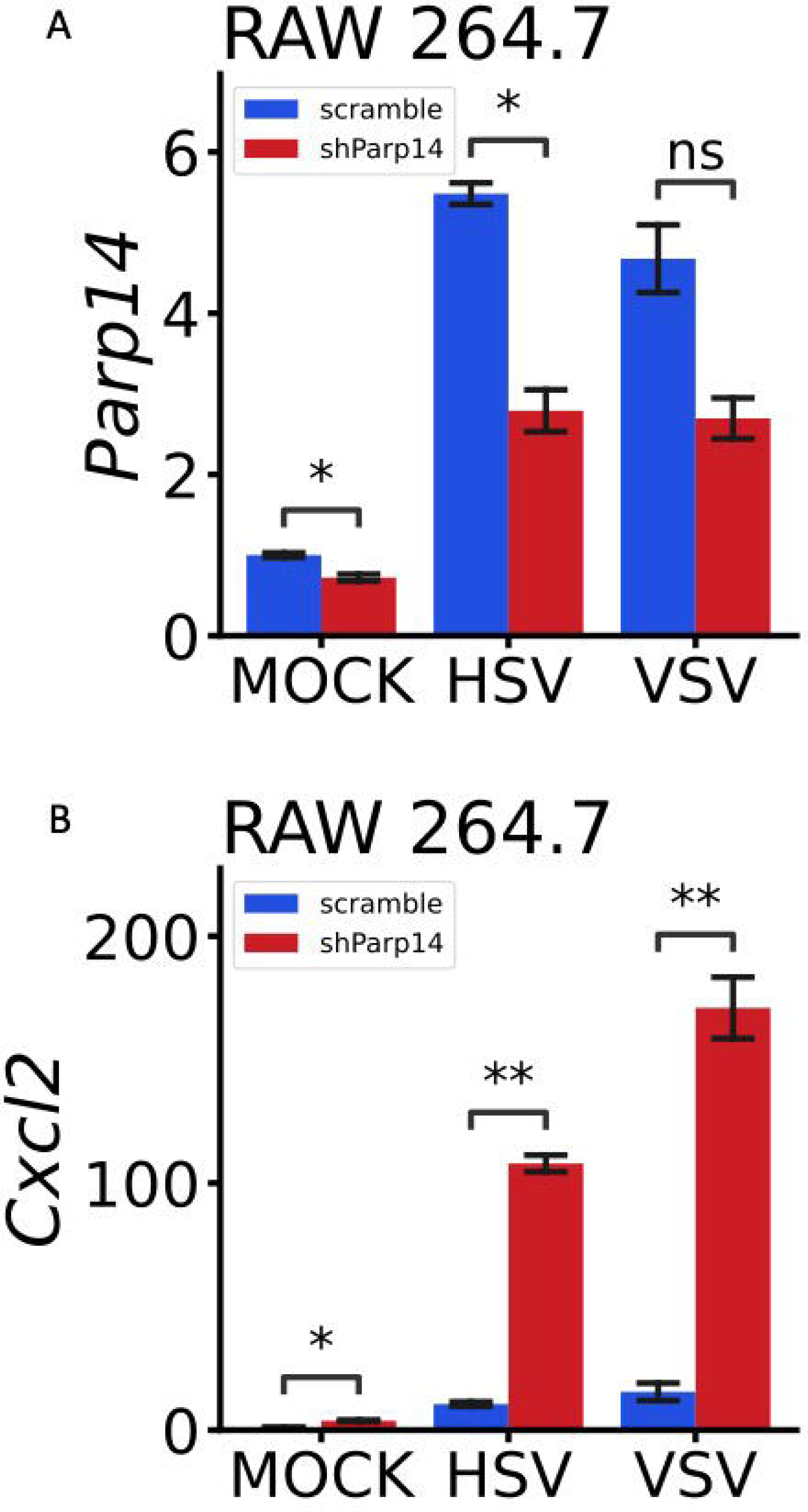
The effect of shRNA for *Parp14*. A) The effect of shRNA to inhibit the expression of *Parp14*. B) Knockdown of *Parp14* could induce the expression of *Cxcl2* significantly.

**Fig. S2.**
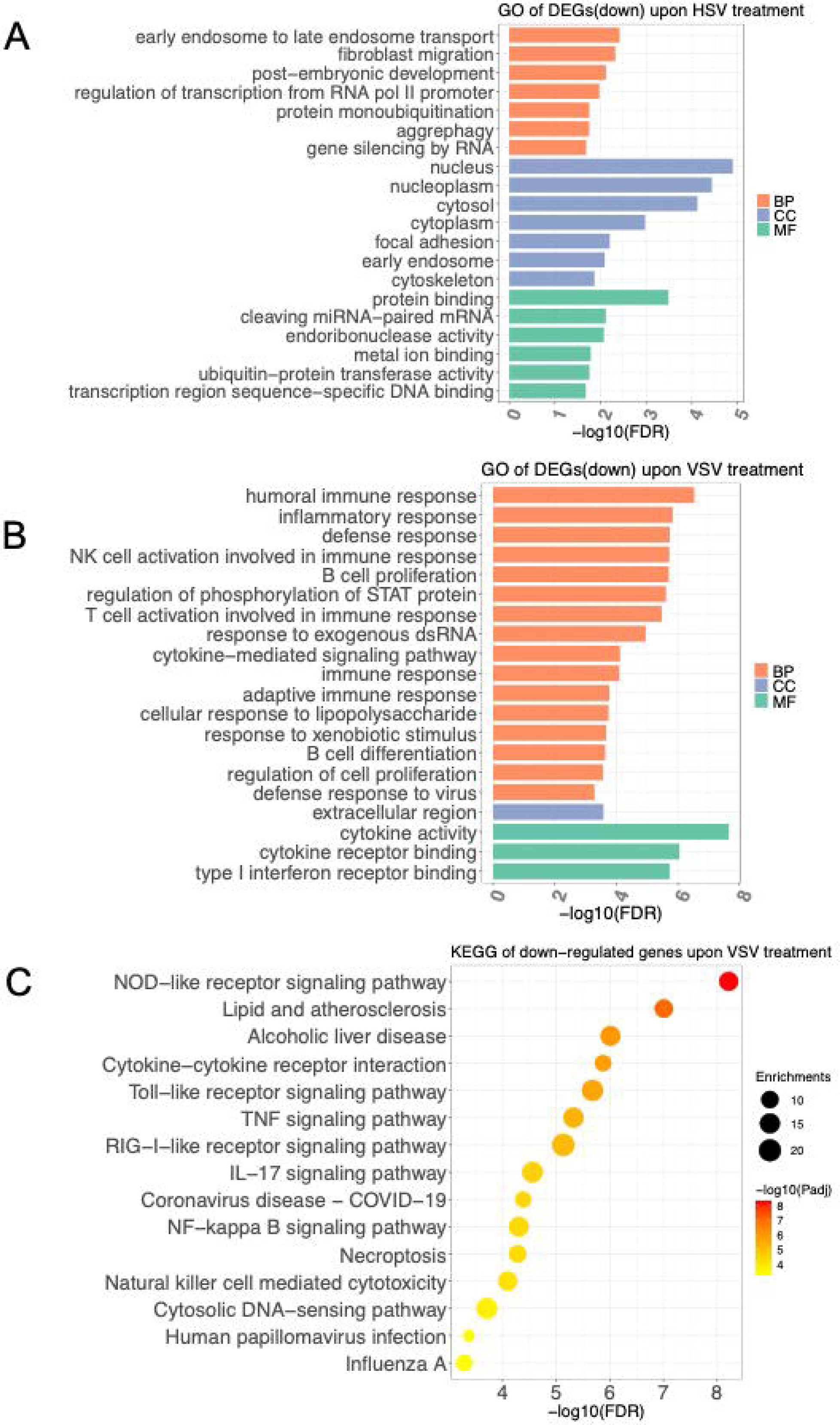
Enrichment analysis of DEGs between *Parp14* KO and WT BMDMs upon HSV and VSV treatment. A) GO annotation of down-regulated DEGs in the *Parp14* KO group compared to WT BMDMs upon HSV treatment. B) GO annotation of down-regulated DEGs in the *Parp14* KO group compared to WT BMDMs upon VSV treatment. C) KEGG pathway enrichment analysis of down-regulated DEGs in the *Parp14* KO group compared to WT upon VSV treatment.

**Fig. S3.**
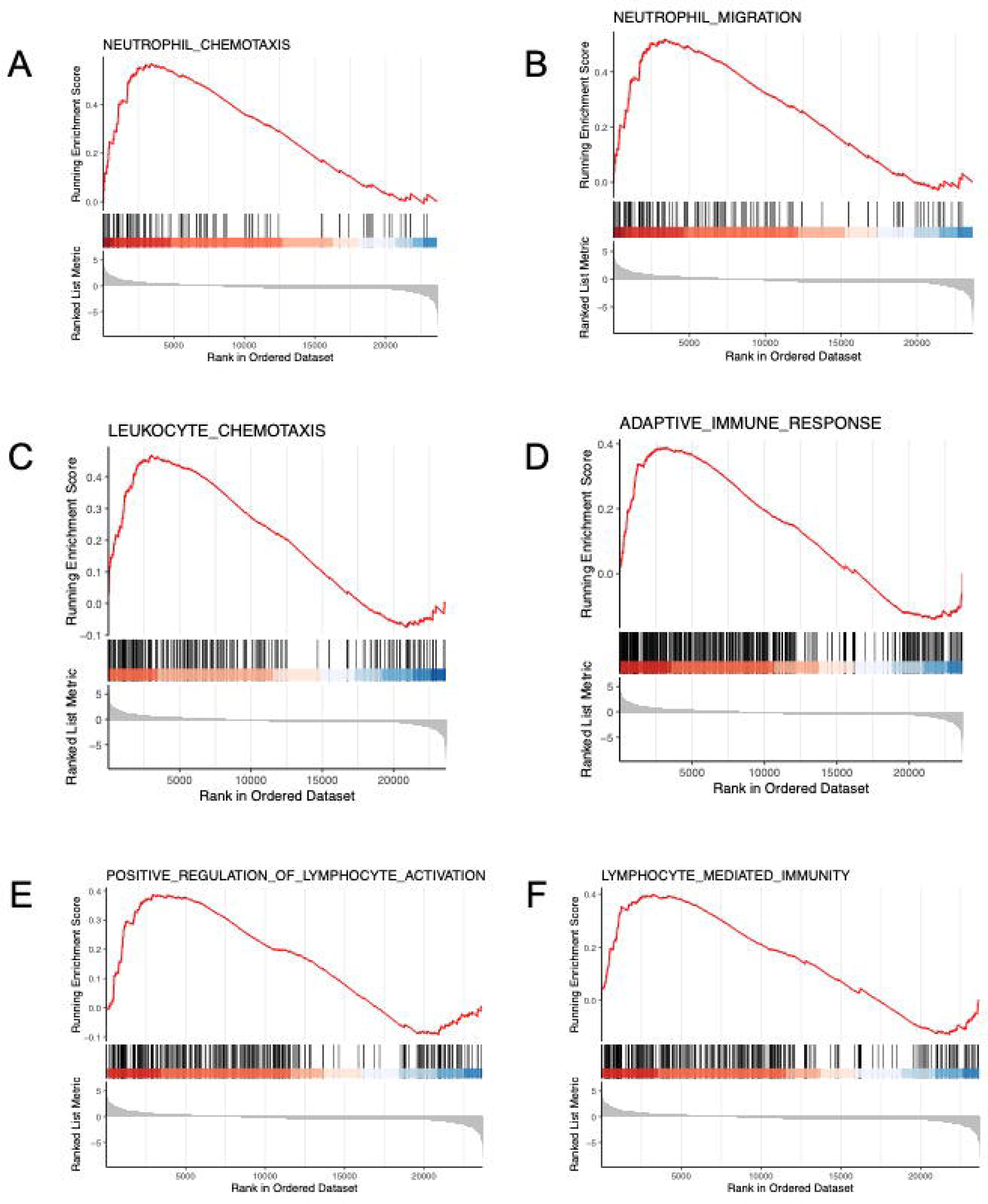
GSEA results of DEGs between the *Parp14* KO and WT PMs upon LPS treatment.

**Fig. S4.**
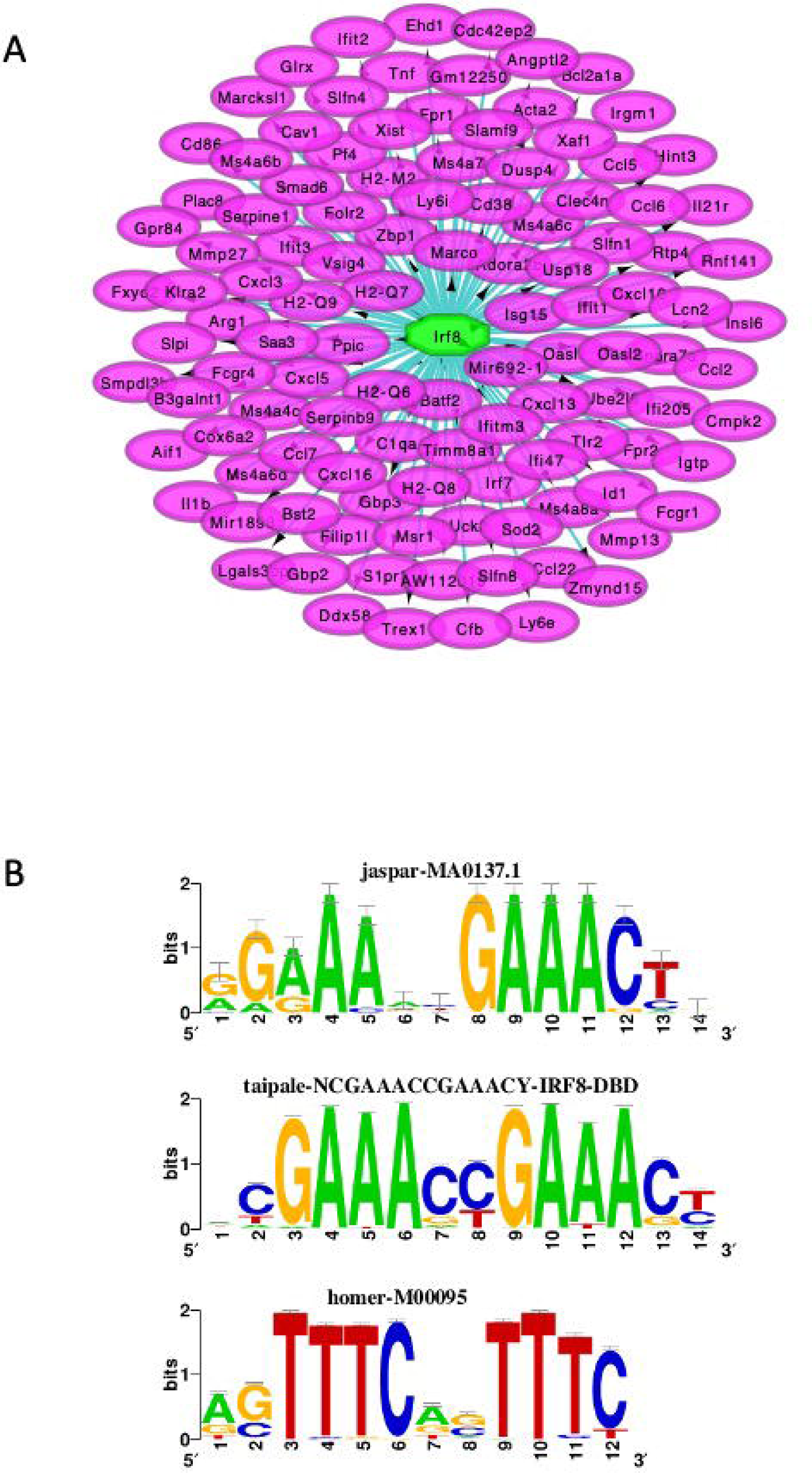
TFEA results of up-regulated DEGs in *Parp14* KO group compared to WT PMs upon LPS treatment. A) IRF8 was the second enriched TF, which may regulate the highly expressed inflammatory cytokines. B) The predicted DNA motifs bound by IRF8.

Table S1. The CRISPR/Cas9 primer sequences to knock out the *Parp14* of mice used in this study.

Table S2. The shRNA primer sequences to knock down the *Parp14* of RAW 264.7 cells used in this study.

Table S3. The primer sequences for RT-qPCR used in this study.

